# WildObserve: Camera-trap software that learns species labels during review and estimates occupancy and abundance

**DOI:** 10.64898/2026.10.06.757089

**Authors:** Michael Tabak

**Affiliations:** Quantitative Science Consulting, LLC, USA; Biology Department, Northern Michigan University, USA

**Keywords:** abundance, camera traps, data privacy, few-shot learning, human-in-the-loop review, object detection, occupancy, species classification

## Abstract

Motion-activated camera traps are an effective tool for monitoring animals, but a single study can produce millions of images, which must be analyzed before they yield data. The burden of analyzing them has led many teams to automate the work with computer vision. These models lose accuracy on images unlike those they were trained on, however, and a general model may not use the categories a project needs. Adapting a model to a project usually requires at least one of the following: many labeled images per species, programming expertise, a graphics processing unit (GPU), or uploading images to a remote server. I describe WildObserve, a desktop application that adapts to each project without any of these requirements. SpeciesNet suggests a species for each animal that MegaDetector finds, and WildObserve keeps SpeciesNet’s feature vector for each one. A project model, a logistic regression on those features, then retrains in about a second as the reviewer labels, learning the project’s own species and names. From the reviewed records, and optionally the suggestions, WildObserve also fits occupancy and abundance models. Images stay on the user’s computer, and every record says which model suggested its species and who confirmed it. WildObserve processed 8.1 images/s on an Apple M1 Pro laptop. On Oregon Critters, a dataset released after SpeciesNet was trained, the project model’s accuracy passed that of SpeciesNet with ten labels per species (balanced accuracy 0.783 against 0.751). I also tested occupancy on 63 cameras in Malawi that SpeciesNet had not seen. With 5% of the images reviewed and the project model’s suggestions counted for the rest, species’ occupancy estimates differed from those of a full review by 0.11 on average. From the reviewed images alone, they differed by 0.41. WildObserve takes a project from images to species records in its own categories and on to occupancy estimates, without writing code, needing a GPU, or sending images elsewhere.

## 1 Introduction

Camera traps have become a standard tool for monitoring wildlife (Burton *et al*., 2015), but a single study can produce millions of images (Norouzzadeh *et al*., 2018). Turning those images into records of which species was where, and when, remains the slowest step. In a global survey, camera-trap users ranked automated filtering of empty images among the most-needed technological developments (Glover-Kapfer *et al*., 2019), and much of the software written for camera-trap projects exists to ease this step (Vélez *et al*., 2023; Hernandez *et al*., 2024).

Machine learning has changed what that step involves. Object detectors such as MegaDetector find animals, people, and vehicles in images from many ecosystems, which lets users set aside the empty images that often make up most of a dataset (Beery *et al*., 2019; Fennell *et al*., 2022; Hernandez *et al*., 2024). Species classifiers trained on large labeled datasets are highly accurate on images like those they were trained on (Norouzzadeh *et al*., 2018; Tabak *et al*., 2019; Willi *et al*., 2019). But accuracy falls when a model meets new locations, species, or camera setups (Beery *et al*., 2018; Schneider *et al*., 2020; Tabak *et al*., 2020; Vélez *et al*., 2023), and a general classifier does not know a specific project’s species list, its naming conventions, or its local look-alike species. Machine labels can substitute for expert labels in ecological analyses once a model has been tested in that ecological context (Whytock *et al*., 2021), but evaluations of AI platforms suggest that most users will still need to review the models’ species predictions (Vélez *et al*., 2023). Projects therefore stand to gain from a model adapted to their own images and categories.

After extracting data from camera trap images, many researchers model species distributions and abundances using models that account for imperfect detection (MacKenzie *et al*., 2002; Royle & Nichols, 2003; Burton *et al*., 2015). Building such models requires converting camera trap data into *detection histories*, each site’s detections by sampling occasion with each camera’s effort, and fitting the models, often in R (R Core Team, 2026) with packages such as camtrapR and unmarked (Niedballa *et al*., 2016; Fiske & Chandler, 2011; Kellner *et al*., 2023). Machine labels complicate this step. Counting images not yet reviewed as non-detections biases occupancy low, whereas counting model labels as detections adds false positives, which bias it high, mostly for rare and look-alike species (Royle & Link, 2006; Miller *et al*., 2011; Whytock *et al*., 2021). Occupancy models that allow for false positives can use confirmed and unconfirmed detections together (Miller *et al*., 2011; Chambert *et al*., 2015; Clare *et al*., 2019), if the records say which detections a person confirmed.

Any model that learns from a reviewer’s labels has to be traceable. As machine labels enter ecological analyses, each record needs to say which model produced a label, how confident the model was, and whether a person checked it (Whytock *et al*., 2021). The Camtrap DP exchange standard has fields for exactly this purpose (Bubnicki *et al*., 2024).

Adapting a model to a project usually requires at least one of four things: many labeled images per species, programming expertise, a graphics processing unit (GPU), or uploading images to a remote server. Many tools now help users manage and label their images (Young *et al*., 2018) and run models and review their output (Vélez *et al*., 2023), and each removes some of these requirements but not all of them. Cloud platforms such as Wildlife Insights host images and classify them centrally (Ahumada *et al*., 2020), so the images must be uploaded. Desktop programs such as Time-lapse (Greenberg *et al*., 2019) and AddaxAI (van Lunteren, 2023) run on the user’s computer, and web applications such as TRAPPER (Bubnicki *et al*., 2016) can be hosted on a project’s own server. MLWIC2 and AddaxAI let users retrain a deep network on their own images without writing code (Tabak *et al*., 2020; van Lunteren, 2023), but retraining still needs many labeled images per species, with accuracy depending strongly on how many (Shahinfar *et al*., 2020), as well as GPUs and choices about network architecture and training. Systems that learn while the user labels cut the labeling effort, by over 99% in one active-learning study (Norouzzadeh *et al*., 2021), but they too retrain deep networks, and AIDE, for example, runs as a web server (Kellenberger *et al*., 2020).

Uploading images is more than an inconvenience. Camera traps photograph people as “human bycatch” (Sandbrook *et al*., 2018), and images and their metadata can reveal the locations of sensitive species, records that need protecting (Lindenmayer & Scheele, 2017; Steenweg *et al*., 2017). Guidance on responsible use asks projects to protect such data and the privacy of the people recorded (Sharma *et al*., 2020; Sandbrook *et al*., 2021). Furthermore, agencies, consultancies, and communities working under data-sharing agreements or privacy law often cannot upload images to a third party. Additionally, teams working with poor internet connections may not be able to upload them at all.

Here I describe WildObserve™, a desktop application that adapts to each project and carries its records through to occupancy and abundance estimates. A model trains in about a second and learns the project’s own species and names as the reviewer labels them. No image is read twice, no GPU is needed, no code is written by the user, and no image leaves the computer. From those records, WildObserve builds detection histories and fits occupancy and abundance models, counting the reviewed images and, if the user chooses, the suggestions on images not yet reviewed. As I describe the software, I ask three questions of it using public datasets. First, how fast does WildObserve process images on an ordinary laptop, and how well do the MegaDetector models it uses find animals? Second, how many labels per species does the project model need to match or pass SpeciesNet’s own suggestions, on cameras it has and has not seen, and how much does the choice of feature extractor matter? Third, how much of a survey must be reviewed before its occupancy estimates come close to those of a full review, with and without suggestions from SpeciesNet or the project model? I answer these questions on datasets that SpeciesNet was probably trained on and others that it has likely never seen.

## 2 Materials and Methods

### 2.1 Design principles

Five principles guided the design. *Local and private*: all processing happens on the user’s computer, and nothing from a project leaves it unless an administrator turns on alerts (described below): no images, thumbnails, or metadata (capture times, locations, camera details, file names), and no annotations, predictions, or accounts. The application sends no telemetry and loads no externally hosted interface assets. It goes online for only four optional tasks, each through a single module that sends only what the task needs: checking for a newer version (sending the application’s version, operating system, and processor type), activating or renewing a license from its key, signing in with Google, and sending alert emails through the organization’s own mail server. An offline mode turns all four off for air-gapped sites; everything else works without an internet connection. The test suite runs with connections to other machines blocked, and tests check what each online request carries and that offline mode sends nothing. *Images stay where they are*: projects reference image folders in place and never modify or share them, unless an administrator turns on email alerts, which can attach the photo. *No machine-learning or programming knowledge required*: users label animals; the model learns from those labels without the user choosing architectures, epochs, or learning rates, and occupancy and abundance models are fitted without writing code. *Traceable*: model output is never edited, human decisions are stored separately, and every change is logged. *Deployable by any organization*: every bundled model and software dependency carries a permissive license, checked automatically on every build, so no component restricts who may use the software or for what purpose.

### 2.2 Architecture

WildObserve is a single program with two parts (Fig. 1). A Python 3.13 back end built on FastAPI stores each project in its own SQLite database and runs every model with ONNX Runtime (ONNX Runtime developers, 2026). ONNX Runtime uses Core ML on Apple computers, CUDA on computers with an NVIDIA graphics card, and the CPU elsewhere, so one model file serves every platform. The interface, written in React and TypeScript, runs in a native window. It talks to the back end only through a server bound to the computer’s loop-back address on a random port. The server rejects requests addressed to any other host name, which blocks web pages from reaching it, and answers only signed-in sessions. People sign in to local accounts with a password or, optionally, with their Google account, in which case Google only confirms their email address; account details stay on the computer. An administrator adds reviewers, and every review is credited to the person signed in. The installers for macOS 14 or later on Apple silicon (signed and notarized), Windows 10 and 11, and Linux bundle the models, so detection and species suggestions work without an internet connection. Large projects stay responsive: on a synthetic project of one million images, a page of the image list loaded in 0.03 s and per-deployment totals in 2 s, because the list is paged and drawn only where it is in view.

**Figure 1.**
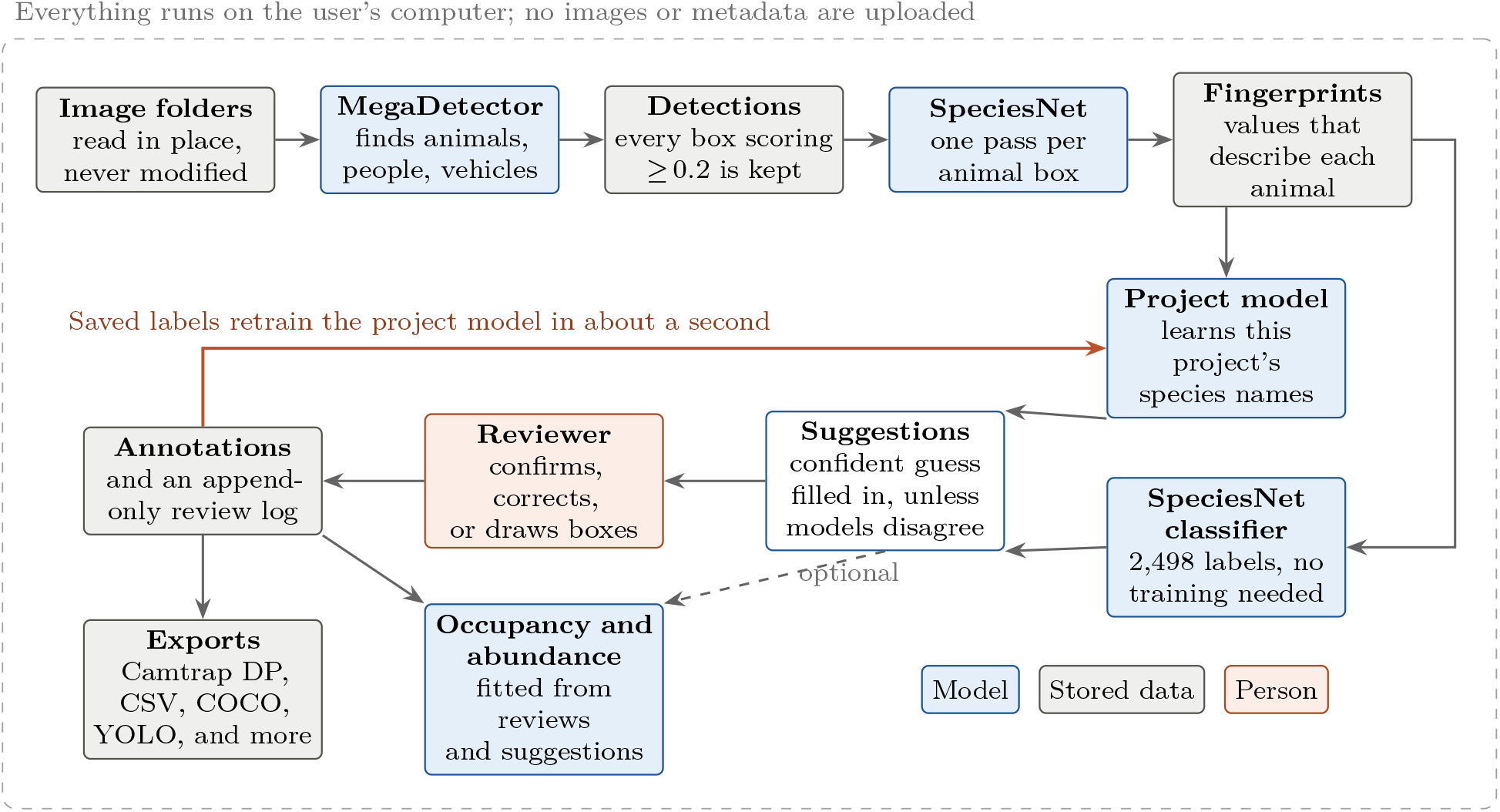
How WildObserve turns camera-trap images into reviewed observations. MegaDetector v6 finds animals, people, and vehicles. SpeciesNet then describes each animal’s appearance as a list of 1,280 numbers, its fingerprint, which is saved so that the network never has to look at that animal again. Two species models read the fingerprints: SpeciesNet’s own classifier, which needs no project labels, and a project model that retrains from the labels the reviewer saves (orange arrow). Model output is never edited; the reviewer’s decisions are stored separately with a full history.

Each project is a folder, by default in `∼/WildObserve/Projects` but wherever the user chooses, such as the drive that holds the images. The folder holds two SQLite databases. The first, `project.db`, contains everything the project produces: the image index, deployments, and capture events; every detection, with the processing run that made it; the reviewer’s annotations and the append-only review log; the fingerprints; and every version of the project model. The second, `thumbs.db`, caches thumbnails and is rebuilt if lost. Copying the folder therefore moves a project, with its annotations and trained project models, to another computer, where its image folders are re-linked; the images themselves never enter the project. Accounts, the license, application settings, and a shared store of imported production models are kept apart from projects, in the application’s own data folder (e.g., `∼/Library/Application Support/WildObserve` on macOS), and the MegaDetector and SpeciesNet models ship inside the application. Because WildObserve keeps project data only on the user’s computer, it makes no copies elsewhere, so back-ups are left to the user. `project.db` holds the work that cannot be recreated (the annotations, review log, and project models), whereas `thumbs.db` is rebuilt if lost. I recommend backing up the whole project folder, for example with the computer’s usual backup software, while WildObserve is closed: while a project is open, the database keeps its latest changes in a separate file beside `project.db`, which a copy of that file alone would miss.

Appendix S1 describes everything WildObserve does, in the order in which a project uses it, and Table S1 lists it: importing and organizing images; detection; species suggestions, from SpeciesNet and the project model; review (Fig. 2); automation, through production models, watched folders, rules, and alerts; records and exports; occupancy and abundance (Fig. 3); and accounts and administration.

**Figure 2.**
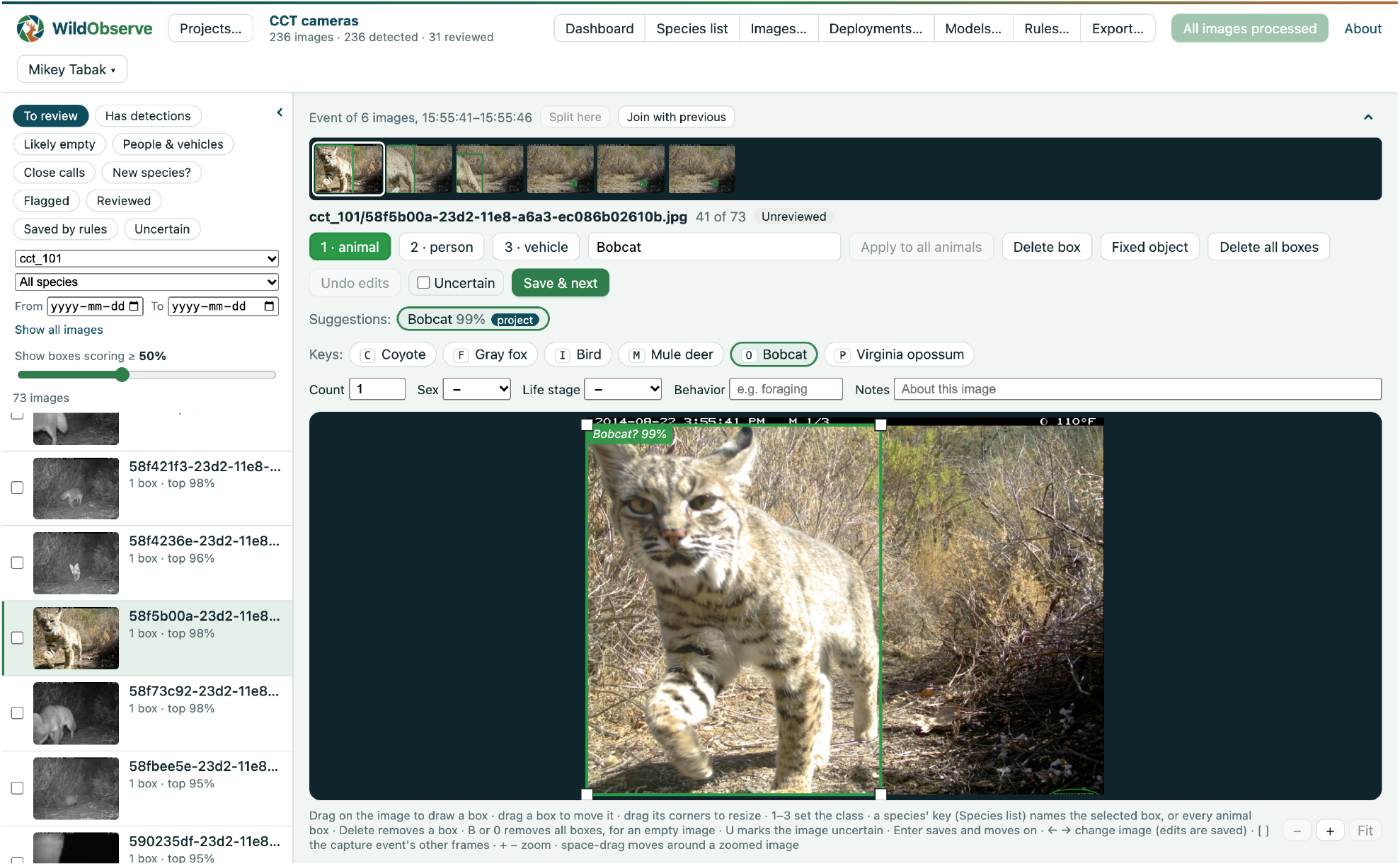
The review screen, here showing images from four Caltech Camera Traps cameras. On the left, the image list shows images to review, with detections, likely empty, with people or vehicles, close calls, new species, flagged, reviewed, saved by rules, or marked uncertain, narrowed by place, camera, deployment, dates, or species; each entry gives its number of boxes and top detection score. The left and top panels can be hidden to more easily view the image. The open image shows a bounding box above the display threshold, which the slider sets (here 50%). Above the image are its capture event, shown as a filmstrip, which can be split or joined, the class buttons, the species field, and the save button; selecting an animal box lists its species suggestions, and the species list’s keys follow. Keyboard shortcuts are listed below the image. Here the list is narrowed to one camera, and the open image, a bobcat, is the first of a six-frame capture event; its suggestion comes from a project model trained on 31 reviewed images.

**Figure 3.**
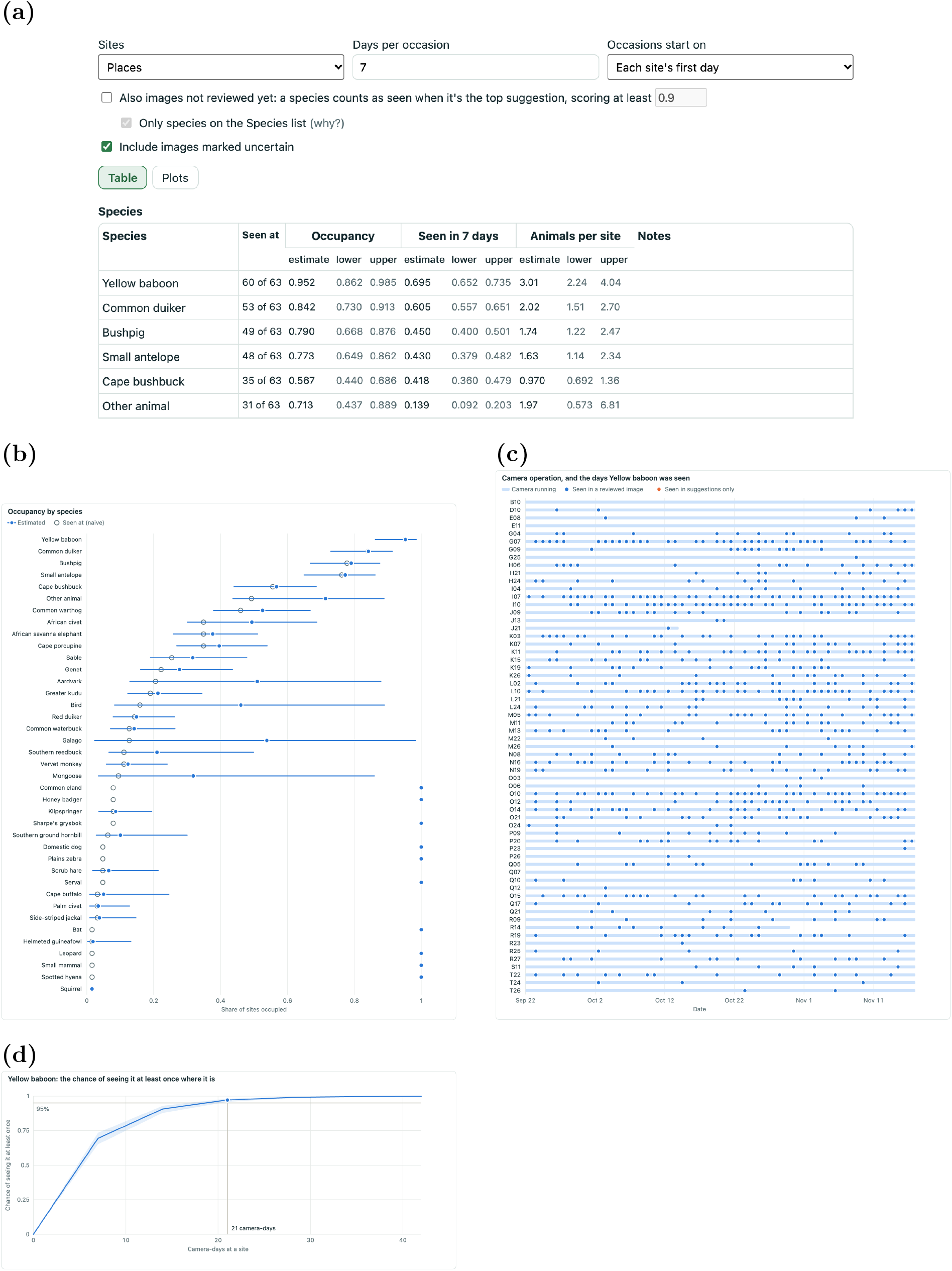
The Occupancy panel, here on the Nkhotakota Camera Traps survey (63 cameras, eight weeks), every image reviewed. (a) Its settings (sites, occasion length, whether suggestions on images not yet reviewed count) and the first rows of its species table: for each species, the sites it was seen at, and occupancy, the chance of detecting it in a seven-day occasion, and Royle–Nichols relative abundance, each with its 95% interval. (b) Occupancy by species, beside the share of sites where each was seen; species seen at too few sites to estimate sit at 1 with no interval, and the table’s notes warn about them. (c) When each camera ran, and the days on which yellow baboons were seen. (d) The chance of seeing yellow baboons at least once at a site they occupy, by camera-days. Each plot can be saved as SVG or PNG.

### 2.3 Software quality

Nearl 300 automated tests cover the review workflow, species training, capture events, exports, accounts, licensing, online requests and offline mode, watched folders, production rules, alerts, occupancy and abundance, and database migrations. The occupancy, Royle– Nichols, and false-positive fits are checked against those of unmarked 1.4.1 (occu, occuRN, and occuFP) on simulated surveys, with and without site covariates, and agree to about four significant figures. Further checks open data saved by each earlier release and compare what it shows, confirm that the bundled models still find the same boxes and species on reference images, and click through the whole interface in a headless browser. Continuous integration runs them on macOS, Linux, and Windows, together with a license check of every locked dependency. A model registry records, for each bundled model, its source, the licenses of its code and weights, its training-data provenance, its SHA-256 checksum, and an approval date. The installer build stops if a bundled model file does not match its registry checksum, and each release records a software bill of materials and the licenses of everything it includes. A smoke test runs each built installer end to end: it signs in, processes images, checks the boxes and species suggestions, and saves a review.

### 2.4 Evaluation data

Evaluations used Caltech Camera Traps (CCT; Beery *et al*., 2018), 243,000 images from 140 camera locations in the southwestern United States, distributed by LILA BC under the CDLA-Permissive-1.0 license. SpeciesNet was trained partly on LILA BC datasets (Gadot *et al*., 2024), probably including CCT, so I repeated the species and detector evaluation on Oregon Critters (Appel *et al*., 2026): 99,909 images from about 1,600 camera locations in western Oregon, United States, from several studies and camera setups, with a box drawn around every animal. LILA BC released it under CDLA-Permissive-1.0 in March 2026, after SpeciesNet’s weights were last changed (versions 4.0.2 and 4.0.3 changed only meta-data) and ten months after the MegaDetector v6 weights were published, so neither model can have been trained on it from LILA BC. Samples were drawn with a fixed random seed, spread across camera locations within each category.

The occupancy evaluation needs surveys in which every image the cameras took was labeled. From CCT, I took every image from the 60 camera locations that ran in the eight weeks from 1 August 2014, the eight weeks in which the most cameras ran (14,956 images). The second survey came from Nkhotakota Camera Traps (Appel *et al*., 2025): 324,635 images from 69 camera locations in Nkhotakota Wildlife Reserve, Malawi, taken from 2018 to 2023 and labeled by image, with counts of animals. LILA BC released it under CDLA-Permissive-1.0 in June 2025, after SpeciesNet’s weights were last changed, and it was labeled in Njobvu-AI rather than Wildlife Insights, so SpeciesNet is unlikely to have been trained on it. I took every image from the eight weeks from 22 September 2020, when the most cameras ran: 41,009 images from 63 cameras, 60 of which ran throughout (58 more, nearly all of people, are not distributed). Both surveys’ images were reduced to 1,024 pixels wide, with each image’s capture time written into its metadata and each camera’s images in a folder of its own, so that each camera formed one deployment and one site.

### 2.5 Detector evaluation

The CCT detector sample contained 650 images from 89 locations: 30 images each of 14 animal categories and of cars, and 200 images labeled empty. The Oregon Critters sample contained 1,140 images of animals, 30 of each of 38 categories, from 680 locations, and 200 images labeled empty; it has no vehicles, and LILA BC removes images of people. I compared the two permissively licensed MegaDetector v6 variants, the compact model (ResNet-18-vd backbone, 80 MB as ONNX) and the large model (ResNet-101-vd, 300 MB), after checking on 40 images that the ONNX conversions reproduced the original PyTorch outputs. Speed was measured on a 2021 MacBook Pro (Apple M1 Pro, 16 GB memory) with ONNX Runtime 1.30, on the CPU alone and with Core ML, using batches of four images and four image-decoding threads, on CCT. On Oregon Critters I measured accuracy only, with Core ML, which gave the same accuracy as the CPU alone on CCT. At thresholds from 0.3 to 0.8 I measured three quantities: *animal recall*, the share of animal images with at least one animal box at or above the threshold; *box recall*, the share of labeled animal boxes matched by a detected box with intersection-over-union of at least 0.5; and the share of empty images with any box at or above the threshold.

### 2.6 Processing speed

To time WildObserve’s processing as a whole, I ran the app’s own processing code on the 650 images of the CCT detector sample, copied into a new project in one folder per camera location and dated by their capture times, so that capture events formed as they had in the field. For each image, WildObserve took its SHA-256 checksum, ran the compact detector, computed SpeciesNet fingerprints for up to five animal boxes, scored them with SpeciesNet’s final layer, and wrote the results to the project, looking for fixed objects and applying the rules after each batch of images. As in the app, the models took one image at a time while two threads read the next images. I then labeled the animal boxes that the detector had found at a score of 0.5 or more with their image’s category (437 boxes in 14 categories), trained the project model, and processed copies of the same images again, each altered by a few bytes so that it was not taken for a duplicate, now scored by both models. I timed each stage, on the CPU alone and with Core ML, on the same laptop.

### 2.7 Species evaluation

I compared models and datasets by their *balanced accuracy*: the share of each species’ boxes that a model names correctly, averaged over species,

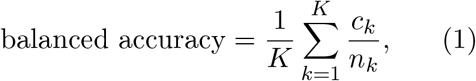

where *K* is the number of species, *n*_*k*_ is the number of boxes of species *k*, and *c*_*k*_ is the number of those that the model names correctly. Averaging over species keeps common species from dominating the score (Brodersen *et al*., 2010), and guessing at random would score 1*/K*.

The species sample contained 1,467 animal crops cut from the labeled boxes of 1,500 images, about 100 in each of 14 categories: bird, bobcat, cat, coyote, deer, dog, fox, lizard, opossum, rabbit, raccoon, rodent, skunk, and squirrel. I compared fingerprints from two frozen feature extractors, each with its standard preprocessing: SpeciesNet (1,280 numbers per crop, as in WildObserve) and DINOv2 ViT-S/14, a general-purpose self-supervised model (384 numbers per crop; square crop with a 10% margin, resized to 224 pixels; Oquab *et al*., 2023). Each was paired with the classifier used in WildObserve. Two scenarios were tested. In the *same cameras* scenario, training and test crops came from the same locations, split by capture sequence so that frames from one burst never fell on both sides; this mimics a project labeling its own images. In the *new cameras* scenario, training crops came only from CCT’s training locations and test crops from its validation locations. For 3, 5, 10, 25, and 50 labels per species, each drawn from a different capture sequence, I trained and tested the classifier on five random draws and report the mean and standard deviation of balanced accuracy (Eq. 1), for which guessing at random would score 0.07 with CCT’s 14 categories and 0.03 with Oregon Critters’ 38. To measure SpeciesNet with no project labels, I summed its probabilities over the labels belonging to each CCT category in SpeciesNet’s taxonomy (for example, every *Urocyon* and *Vulpes* label counted as fox) and computed balanced accuracy on all 1,467 crops.

The Oregon Critters sample contained 3,831 crops from 3,677 images in 38 categories (23 of mammals, 14 of birds, and one of reptiles and amphibians), with up to 100 images each; Canada jay, marten, and coyote had fewer. Its categories mix species with groups such as “*Catharus* species”, “weasel family”, and “other bird”. Its published split is by image rather than by location, so for the new-cameras scenario I held out a random quarter of its camera locations, and I treated a location’s images taken less than a minute apart as one capture sequence. Each SpeciesNet label counted toward the narrowest category containing its taxon; a broader label counted only when a single category lay within it, so labels such as “Sciuridae family” counted as wrong. SpeciesNet often names a small bird only as “Bird”, where Oregon Critters names its species or family. I counted that label as correct for any bird, as CCT’s single bird category did implicitly, and also report the result without this allowance.

### 2.8 Occupancy evaluation

I processed each survey with WildObserve and, on copies of the project, saved reviews from the dataset’s own labels as a reviewer would. On an image of one species, every animal box the detector found at 0.5 or more was named with that species, and the image’s count of animals was spread over them. An image without animals was saved as empty. An image showing several species, or a species on which the detector put no box, received a box over the whole frame for each species and was marked uncertain, which keeps it out of the project model’s training but not out of the occupancy tables. Saving every image this way gave a full review, the standard that the other estimates were compared with.

I then reviewed a share of each survey’s images (2, 5, 10, 25, and 50%), choosing whole capture events at random until the share was reached, five times for each share, with a fixed random seed for each share. From each partial review I estimated occupancy and abundance three ways: from the reviewed images alone; with SpeciesNet’s suggestions on the other images, without a project model; and with suggestions from a project model trained on the reviewed images. With suggestions, I estimated occupancy both with the standard model and allowing for false positive detections (Appendix S1, Eq. S2). I also estimated occupancy from SpeciesNet’s suggestions with no review at all. WildObserve’s defaults applied throughout: places as sites, seven-day occasions from each site’s first day, detector boxes from 0.5, suggestions scoring at least 0.9, and only species on the project’s list, which held the dataset’s categories. SpeciesNet’s labels counted toward each dataset’s categories by taxonomy, as in the species evaluation; categories without a SpeciesNet label, such as Nkhotakota’s “small antelope”, were scored only with the project model. On Nkhotakota, each camera’s set-up and take-down times came from the first and last images on its memory cards.

For each species, I compared occupancy estimated from each partial review with occupancy estimated from the full review. I scored only the species that the full review estimated well: those it detected at three or more sites, whose model fit converged without reaching a limit (an occupancy of 0 or 1), and whose 95% interval for occupancy was no wider than 0.5, since a less precise estimate gives no firm value to compare with. For each way of estimating occupancy, I report the mean, over the scored species, of the absolute difference between a species’ occupancy and its occupancy from the full review (occupancy runs from 0 to 1, so a difference of 0.1 is 10 percentage points); the mean signed difference (bias); and the share of 95% intervals that held the full review’s estimate, each averaged over the five draws. A species that a run never detected counted as occupancy 0. Where no reviewed image showed a species, the false-positive model gives no estimate, and the standard estimate stood in. I also counted false detections, occasions (a site’s seven-day periods) in which a run detected a species that the full review did not, and missed detections, the reverse, and compared Royle–Nichols abundance by the median ratio of each estimate to the full review’s.

## 3 Results

### 3.1 Detection

The ONNX conversions matched the original models, giving identical detections on all 40 test images; scores differed by at most 5*×*10^−6^ (compact) and 5 *×* 10^−4^ (large). The compact model processed 4.9 images/s on the CPU, about 420,000 images a day, and 38.5 images/s with Core ML (Table 1). The large model was about four times slower on the CPU and three times slower with Core ML. Core ML compiled the RT-DETR models only when the batch size was fixed in advance, and the first load took 23 s (compact) and 51 s (large). Decoding JPEG images at reduced resolution halved decoding time (9.6 vs. 18.4 ms per image) and gave the same result at a 0.2 threshold on 98 of 100 images.

**Table 1.** Speed on the 650 Caltech Camera Traps images, on a 2021 MacBook Pro (Apple M1 Pro, 16 GB), with ONNX Runtime 1.30. *MegaDetector alone*: the two permissively licensed MegaDetector v6 variants (backbone, file size), in batches of four images with four decoding threads. *WildObserve processing*: the app’s own processing, one image at a time with two threads reading ahead: each image’s checksum, the compact detector, SpeciesNet fingerprints of its animal boxes (2.5 per image), species scores, and writing the results to the project, in a new project and again with a trained project model. *ms/image*: model time only. *images/s*: end to end, including reading and decoding the images. *First load*: time to compile the models for Core ML the first time they are used.

|  | CPU only |  | Core ML |  |  |
| --- | --- | --- | --- | --- | --- |
|  | ms/image | images/s | ms/image | images/s | first load (s) |
| *MegaDetector alone* |  |  |  |  |  |
| Compact (ResNet-18-vd, 80 MB) | 199 | 4.9 | 29 | 38.5 | 23 |
| Large (ResNet-101-vd, 300 MB) | 757 | 1.3 | 97 | 11.6 | 51 |
| *WildObserve processing (compact detector and SpeciesNet)* |  |  |  |  |  |
| New project | 666 | 1.5 | 116 | 8.1 | 24 |
| With a project model | 672 | 1.5 | 114 | 8.1 | – |

The two variants found animal images about equally well; the large model found more individual boxes, especially at high thresholds (Table 2). Both found every car at thresholds up to 0.8. At the default 0.5 threshold, the compact model found an animal in 97.1% of animal images and matched 88.8% of labeled animal boxes, but also put a box on 48.0% of images labeled empty. At 0.7 that share fell to 12.5%, while animal recall fell to 87.4%. Inspection of the highest-scoring boxes on empty images showed two causes. Some were label errors in the CCT dataset: the highest-scoring box, at 0.97, was a person standing in an image incorrectly labeled empty. Most others were small boxes (median 0.8% of the image) on static objects such as stumps, concentrated at a few cameras: three locations produced 49 of the 96 images flagged at 0.5. The median top score on empty images was 0.49 (compact) and 0.52 (large), well above the 0.2 threshold often used with MegaDetector. Storing boxes down to 0.2 kept at most 18 boxes per image (median 2), whereas a floor of 0.05 would have kept up to 293.

**Table 2.**
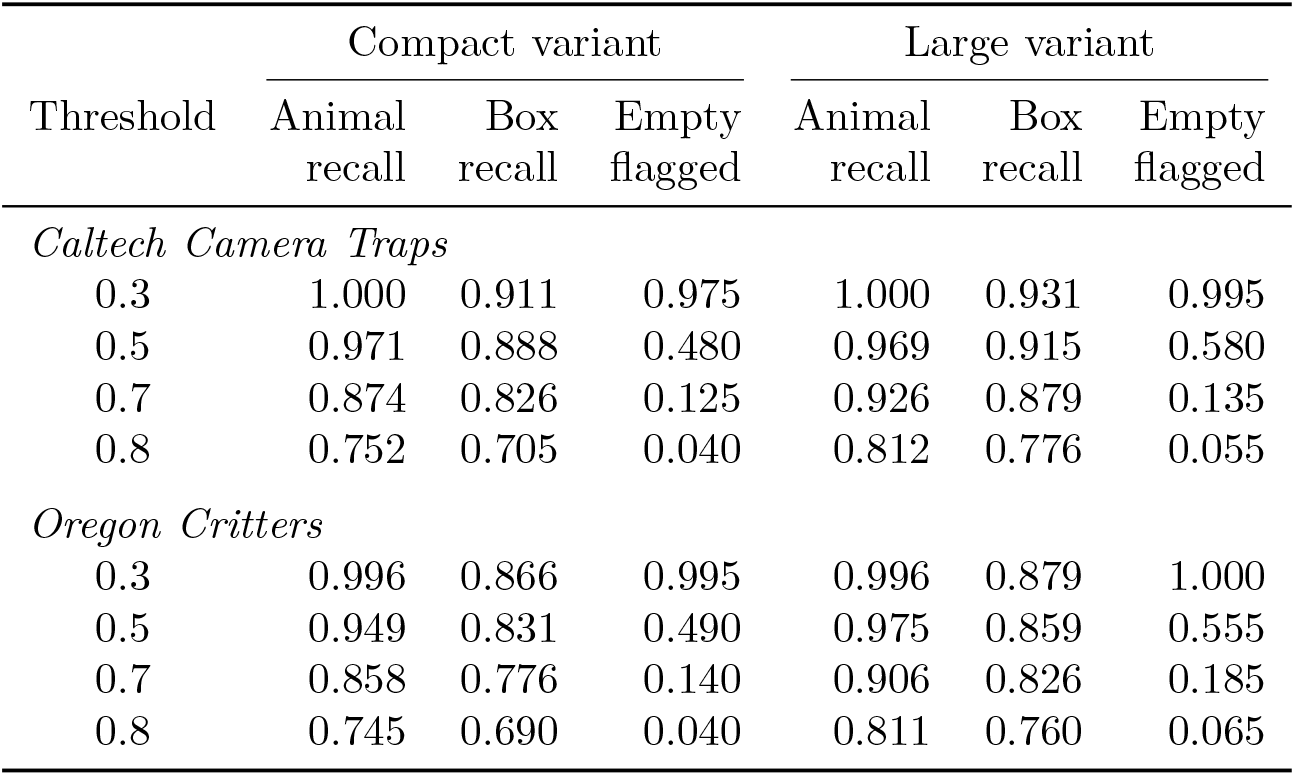
Detection accuracy of the two MegaDetector v6 variants on 650 Caltech Camera Traps images (450 with animals or cars, 200 labeled empty) and 1,340 Oregon Critters images (1,140 with animals, 200 labeled empty), by confidence threshold. *Animal recall*: share of animal images with at least one animal box at or above the threshold. *Box recall*: share of labeled animal boxes matched at intersection-over-union ≥ 0.5. *Empty flagged*: share of empty images with any box at or above the threshold; some of CCT’s empty images contain people or animals that the labels missed.

On Oregon Critters, which neither model had seen, both matched fewer of the labeled boxes than on CCT, and the compact model found animals in fewer images (Table 2). At 0.5, the compact model found an animal in 94.9% of animal images and matched 83.1% of labeled boxes, and the large model 97.5% and 85.9%; where the two models had found animals equally well on CCT at this threshold, here the large one led. Boxes on empty images were as common as on CCT: at 0.5, 49.0% of empty images (compact) and 55.5% (large) had one, and at 0.7, 14.0% and 18.5%. The median top score on empty images was again about 0.5 (0.50 compact, 0.52 large).

### 3.2 Processing speed

WildObserve processed the 650 CCT images at 8.1 images/s with Core ML, about 700,000 images a day, and at 1.5 images/s on the CPU alone, about 130,000 a day (Table 1). Fingerprinting took most of the time: WildObserve fingerprinted 2.5 animal boxes per image on average (every animal box scoring 0.2 or more, up to five), at 34 ms each with Core ML and 186 ms on the CPU. Of the 124 ms each image took with Core ML, finger-printing took 86 ms, detection 30 ms, and the checksum, species scores, and database 9 ms, while another thread spent 21 ms reading the next image. On the CPU, fingerprinting took 462 ms and detection 204 ms of 675 ms. A trained project model did not slow processing (8.1 images/s with Core ML, 1.5 images/s on the CPU). Training it on the 437 labeled boxes took 0.36 s, and re-scoring all 1,616 fingerprinted boxes with it took 0.65 s. Loading both models took 5 s once they had been compiled for Core ML, and 24 s the first time.

### 3.3 Species suggestions

With no project labels, SpeciesNet reached 0.919 balanced accuracy on the 14 categories; on average, 83% of its probability fell on labels that belong to one of them. A project model on SpeciesNet fingerprints reached 0.886 *±* 0.017 with three labels per species, nearly matched SpeciesNet alone with five labels (0.914 *±* 0.005), and leveled off at 0.93–0.94 from 25 labels (Fig. 4). Balanced accuracy on new cameras was within 0.016 of that on the same cameras at every label count (0.883 vs. 0.886 at three labels; 0.931 vs. 0.937 at 25). Fingerprints from DINOv2 were much weaker: 0.578 with three labels and 0.784 with 50 on the same cameras, and 0.740 with 50 on new cameras. The ONNX conversion of SpeciesNet gave the same top label as the original model on all 20 crops tested.

**Figure 4.**
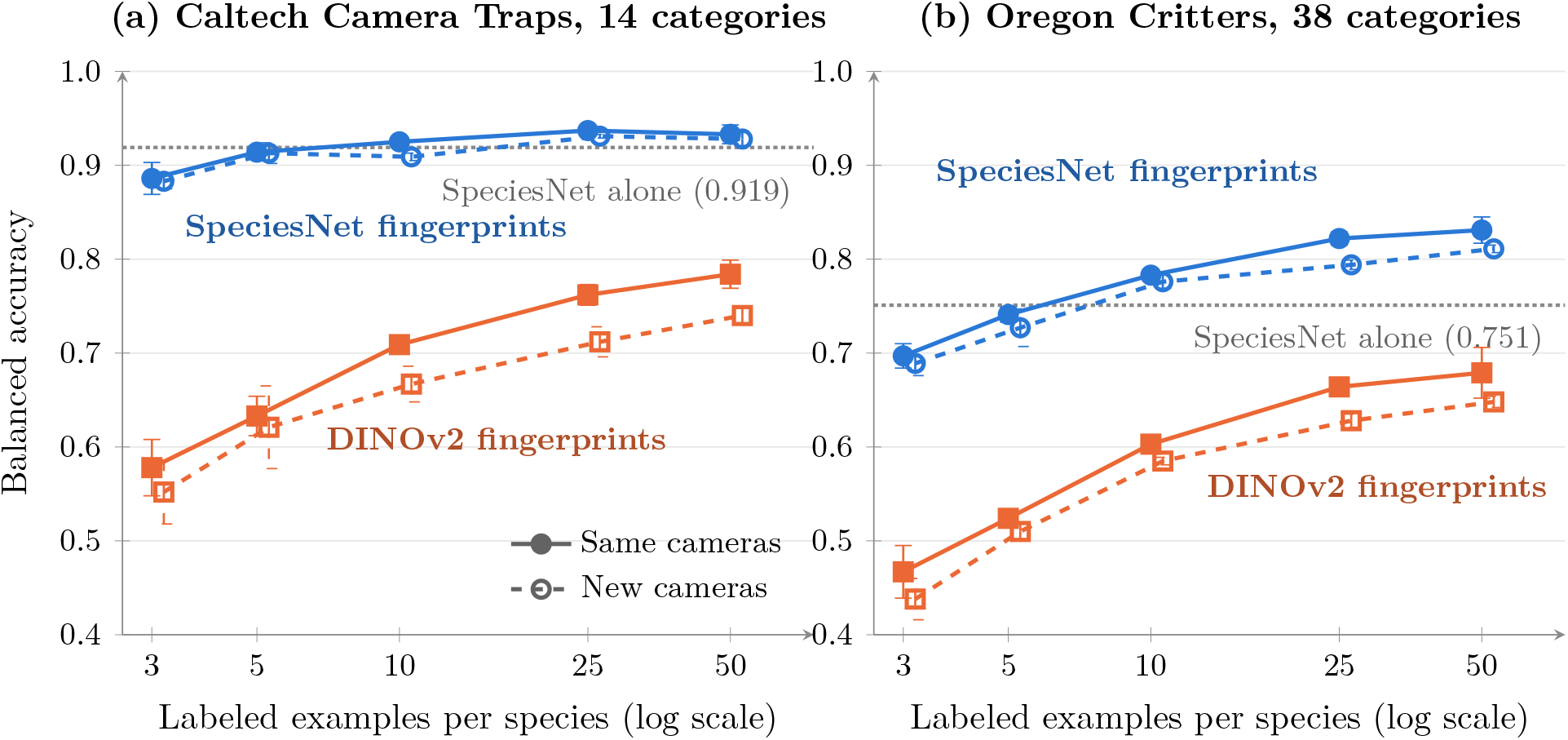
Species accuracy of a project model by the number of labeled examples per species, (a) on 1,467 Caltech Camera Traps crops in 14 categories, a dataset SpeciesNet was probably trained on, and (b) on 3,831 Oregon Critters crops in 38 categories, which SpeciesNet had not seen. Each point is the mean balanced accuracy over five random draws of training labels; bars show *±*1 standard deviation. Blue: fingerprints from SpeciesNet, as used in WildObserve; orange: fingerprints from DINOv2 ViT-S/14, a general-purpose model. Solid lines: training and test images from the same cameras; dashed lines: test images from camera locations absent from training. The dotted line shows SpeciesNet’s own classifier with no project labels; in (b) its “Bird” label counts as correct for any bird category (0.528 without that allowance).

On Oregon Critters, every balanced accuracy was lower (Fig. 4b). SpeciesNet alone reached 0.751 balanced accuracy on the 38 categories, or 0.528 without the allowance for “Bird”; on average, 69% of its probability fell on labels that belong to a category. It stayed accurate on medium and large mammals: 0.88 on the 11 categories that fall within CCT’s sampled ones (bobcat, deer, elk, gray fox, dog, coyote, opossum, raccoon, two skunks, and rabbits), 0.95 on black bear, and 0.87 on cougar. It did worst on the region’s smaller and less familiar animals: mountain beaver (0.00, most often called “Mammal”), woodrats (0.07, “Rodent”), reptiles and amphibians (0.16, mostly “Blank”), and martens and weasels (about 0.54). For 60% of bird crops its best answer was “Bird”, and for 20% the correct species or family. The project model on SpeciesNet fingerprints reached 0.697 *±* 0.013 with three labels per species, passed SpeciesNet alone at ten (0.783 *±* 0.006), and reached 0.831 *±* 0.014 with 50. On new cameras it reached 0.689, 0.776, and 0.811, at most 0.028 below the same cameras. DINOv2 fingerprints reached 0.467 with three labels and 0.679 with 50 (0.648 on new cameras), 0.15 to 0.25 below SpeciesNet’s.

### 3.4 Occupancy

On the Nkhotakota dataset, 20 of the 39 species detected in the full review were scored (Fig. 5a). With SpeciesNet’s suggestions alone and no review, a species’ occupancy differed from its occupancy in the full review by 0.242 on average (median 0.108). SpeciesNet’s suggestions detected a species in only 9 occasions in which the full review had not, but missed it in 814 in which the full review had. Much of the difference came from names: SpeciesNet called the dataset’s yellow baboons, seen at 60 of the 63 sites, “Kinda baboon”, a related species, on 3,999 boxes scoring 0.9 or more; it called bushpigs “wild boar” and many duikers and other antelopes “white-tailed deer”. A project that names the species it has does not count these suggestions. Counting SpeciesNet’s suggestions beside a partial review narrowed the difference little until a quarter of the images were reviewed (0.233 with 2% reviewed, 0.156 with 25%, and 0.078 with 50%). The project model’s suggestions came much closer: 0.182 with 2% of the images reviewed, 0.107 with 5%, 0.083 with 10%, and 0.048 with 50%, against 0.458, 0.410, 0.382, and 0.094 for the reviewed images alone (Fig. 5a). The project model’s suggestions detected species in more occasions in which the full review had not (177 to 239), but missed fewer (81 to 400), and from 5% reviewed its 95% intervals held the full review’s estimate for 94 to 96% of species. Its relative abundance was 7 to 20% above the full review’s.

**Figure 5.**
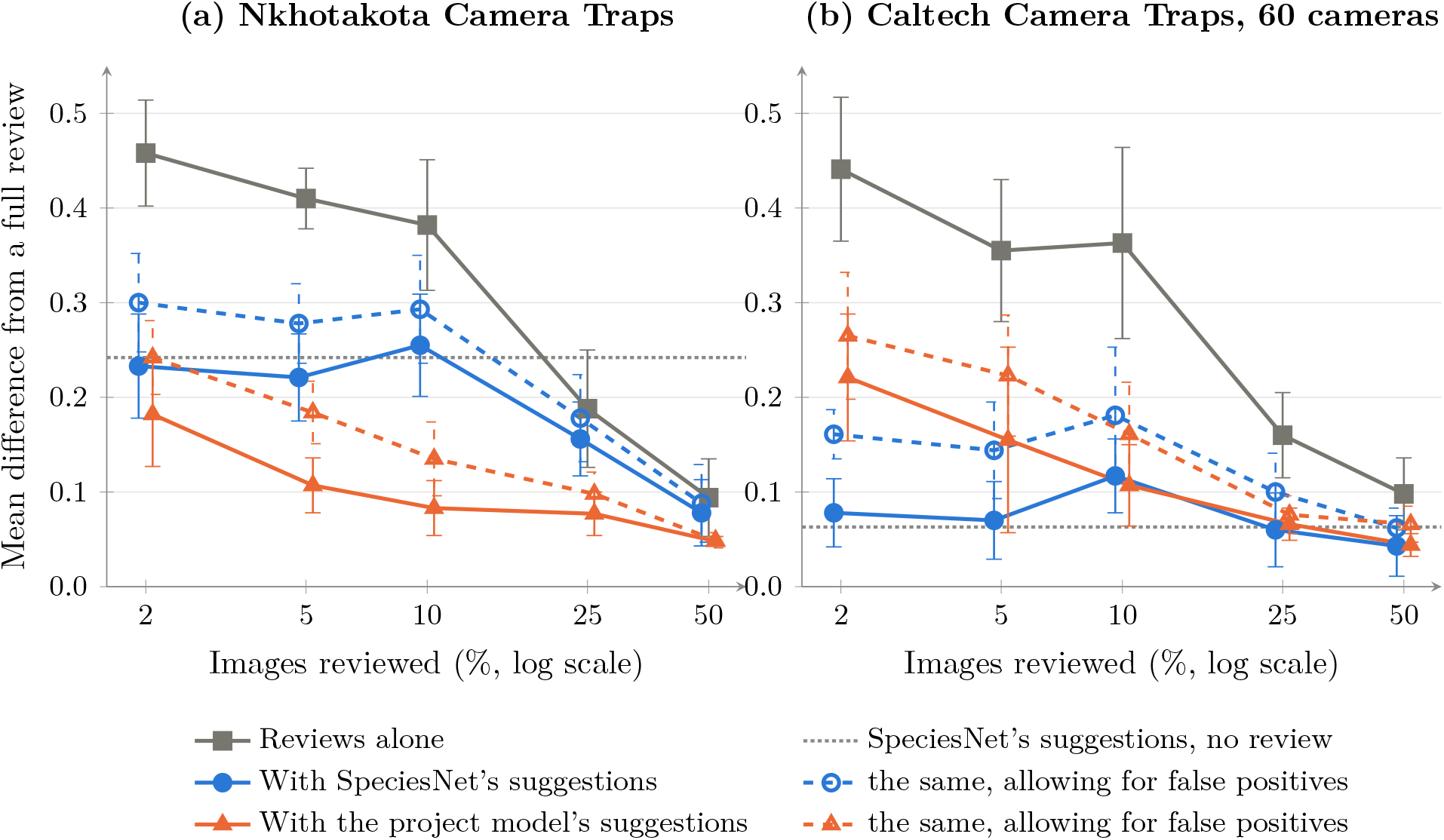
How far occupancy estimated from part of a survey’s review falls from a full review’s, by the share of images reviewed, (a) on Nkhotakota Camera Traps, which SpeciesNet had not seen, and (b) on 60 cameras of Caltech Camera Traps, which it probably had. Each point is the mean, over the species scored, of the absolute difference between a species’ occupancy and the full review’s, averaged over five random draws of capture events to review; bars show *±*1 standard deviation over the draws. Grey: reviewed images alone. Blue: with SpeciesNet’s suggestions on the images not reviewed; orange: with the project model’s, trained on the reviewed images. Dashed lines: allowing for false positives among the suggestions. The dotted line shows SpeciesNet’s suggestions with no review at all. Species are scored when the full review detected them at three or more sites and its 95% interval for their occupancy was no wider than 0.5 (20 of 39 species on Nkhotakota, 10 of 16 on CCT).

On CCT, 10 of the 16 species detected in the full review were scored (Fig. 5b). With SpeciesNet’s suggestions alone, a species’ occupancy differed from the full review’s by 0.063 on average (median 0.029), and 89% of the 95% intervals held the full review’s estimate; their relative abundance was within 1% of the full review’s (median ratio 0.99). With a partial review as well, the difference stayed between 0.043 and 0.078, except with 10% reviewed (0.117). There, in four of the five draws, the suggestions found lizards at only one or two sites, and the estimate of their occupancy reached its upper limit of 1. WildObserve’s Occupancy panel marks such an estimate with warnings that the species was seen at too few sites and that its occupancy is at its limit. Reviewed images alone again gave estimates far from the full review’s until much of the survey was reviewed (0.441 with 2%, 0.160 with 25%, and 0.098 with 50%). The project model’s suggestions were less accurate than SpeciesNet’s with little review (0.221 against 0.078 with 2% reviewed) and as accurate with half the survey reviewed (0.044 against 0.043). The project model’s suggestions overestimated occupancy, by 0.038 to 0.098 on average, because they added detections that the full review did not have, in 77 to 87 occasions, birds more often than any other species.

On both surveys, allowing for false positives made the estimates worse and biased them low, except with the project model at 50% reviewed on Nkhotakota, where it changed little (0.049 against 0.048). Counting only the project model’s suggestions that SpeciesNet agreed with, as the review fills them in, cut the false detections by half or more on both surveys. It brought occupancy closer to the full review’s on CCT (0.017 rather than 0.044 with 50% reviewed) but not on Nkhotakota (0.051 rather than 0.048), where it missed more. Scoring every species, including those whose full-review interval was wider than 0.5, changed the means by at most 0.06 (Fig. 5).

## 4 Discussion

WildObserve pairs a general species classifier with a project model that learns from a handful of labels. On Caltech Camera Traps, three labels per species gave suggestions almost as accurate as SpeciesNet’s own, about five matched it, and 25 slightly exceeded it, in the project’s own species names. On Oregon Critters, which SpeciesNet had not seen, about ten labels per species passed SpeciesNet’s own suggestions. Because the model learns from stored fingerprints, it trains in about a second on a laptop CPU, and re-scoring every box in a project is a single matrix multiplication. Users therefore see better suggestions after each save, without reprocessing images, without a GPU, and without programming or conceptualizing a computer vision model. Retraining a deep network, the usual way to adapt a classifier to a project (Tabak *et al*., 2020; van Lunteren, 2023), needs far more labels (Shahinfar *et al*., 2020), a GPU, choices about how to train the network, and often computer programming expertise.

The results support earlier work on few-shot learning, where a strong pretrained representation and a linear classifier rival more elaborate methods (Chen *et al*., 2019; Tian *et al*., 2020), and the finding that better pretrained networks transfer better (Kornblith *et al*., 2019). The choice of feature extractor mattered more than any other factor: fingerprints from SpeciesNet, trained on cameratrap images, beat those from a general-purpose model by 0.15 in balanced accuracy even with 50 labels per species, on both datasets. On CCT, SpeciesNet fingerprints also scored at most 0.016 lower on new camera locations, whereas DINOv2’s scored up to 0.050 lower; on Oregon Critters, at most 0.028 and 0.036 lower. Classifiers trained on narrower data often generalize poorly to new locations (Beery *et al*., 2018; Schneider *et al*., 2020); a feature extractor trained on tens of millions of images from many cameras appears to have learned features that carry over.

If SpeciesNet alone scores 0.92 on CCT, why add a project model? Oregon Critters gives the plainest answer: on images it had not seen, SpeciesNet’s balanced accuracy was 0.75, and was near zero for some of the region’s small mammals, while ten labels per species gave the project model a higher balanced accuracy than SpeciesNet. First, a project model speaks the project’s language. It uses the project’s species list and names, and it can learn project-relevant groupings (e.g., “small rodent”) or species that SpeciesNet lacks or often assigns only to a higher taxon. Second, it chooses only among the project’s species. SpeciesNet spreads its probability over thousands of labels (17% of it fell on labels outside CCT’s 14 categories, and 31% outside Oregon Critters’ 38), which is probably why the project model leveled off slightly above SpeciesNet alone on CCT, on new cameras as well as familiar ones. Third, the two models check each other. A project model knows only the species it has been taught and can be confidently wrong about anything else; SpeciesNet knows thousands of labels but not the project’s names. Filling in a suggestion only when both agree, and flagging the box when they do not, keeps the fast path for routine images and sends unusual ones to a closer look.

With species suggestions included, WildObserve processed about 130,000 images a day on a laptop’s CPU alone and 700,000 with Core ML, enough for a large archive on an ordinary laptop. Fingerprinting the animal boxes, not detection, took most of that time, and because the project model reuses the stored fingerprints, it added no measurable time. The detector results also bear on how thresholds should be set. On both datasets, its scores on empty images were high enough that a threshold suited to earlier MegaDetector versions would put a bounding box on nearly every empty image. Because WildObserve stores every box above 0.2 and applies the display threshold when data are read, users can tune the threshold to their own images later. The boxes on static objects that recur at particular cameras called for a dedicated tool: WildObserve finds boxes that appear in the same place across many capture events from one camera and, once a reviewer confirms them, hides these fixed objects everywhere.

WildObserve also takes a project past its records to occupancy and abundance, the estimates most camera-trap studies are run for (Burton *et al*., 2015), without exporting data or writing code. The occupancy evaluation shows how SpeciesNet and the project model fit into that step. On CCT, which SpeciesNet had probably seen, its suggestions alone gave occupancy within about 0.06 of a full review’s, whereas reviewed images alone needed about half the survey reviewed to come within 0.1 (Fig. 5): images not yet reviewed count as non-detections, which lowers the apparent chance of detection and makes occupancy hard to separate from it. On Nkhotakota, which SpeciesNet had not seen, its suggestions were much less useful (a mean difference of 0.24 from the full review’s occupancy), mostly because it named several of the reserve’s species as related species from elsewhere, names a project would not use. The project model, which learns the project’s own names from its reviews, came within about 0.1 of the full review with 5 to 10% of the images reviewed, where reviewed images alone were about 0.4 away. This is the case for a project model in occupancy work: a modest review teaches it the project’s species, and its suggestions then stand in for the rest. The surveys also differ in their categories. CCT’s are broad (any deer is “deer”), which forgives the species-level confusions that cost SpeciesNet on Nkhotakota, so part of the gap between the surveys comes from the categories rather than from what SpeciesNet had seen.

The project model’s suggestions also added false detections. A model that knows only the project’s species gives every box one of them, including boxes on empty frames, and on CCT it did so most often for birds. The review guards against this by filling in a project-model suggestion only when SpeciesNet agrees. Applying the same rule to the occupancy tables cut false detections by half or more, but improved occupancy only on CCT; on Nkhotakota, where SpeciesNet’s names often differ from the project’s, it discarded true detections too, so the tables count every confident suggestion. Allowing for false positives (Miller *et al*., 2011) did not help in these surveys: it made estimates worse and biased them low. SpeciesNet’s suggestions had few false positives to correct, and although the project model’s had many, a partial review confirmed too few detections to tell a species that is present but seldom confirmed from a false positive, which such models need (Chambert *et al*., 2015; Clare *et al*., 2019). They may still be worth their wider intervals where false positives are common, as with look-alike species, and many detections are reviewed.

Keeping images and records on the user’s computer addresses the concerns about people in camera-trap images (Sandbrook *et al*., 2018; Sharma *et al*., 2020) and about disclosing sensitive locations (Lindenmayer & Scheele, 2017). It also means no upload time, no cloud costs, no dependence on an internet connection in the field, and no reliance on data centers. The trade-off is that responsibility for security, backups, and access control stays with the user. Keeping model output and human decisions apart, logging every change, and recording the model and score behind every accepted suggestion lets analyses separate verified from unverified records (Whytock *et al*., 2021). The Camtrap DP export fills the standard’s fields for how, by whom, and when each observation was classified (Bubnicki *et al*., 2024).

These evaluations have clear limits. Speed was measured on CCT alone, and it depends on how many animals each image holds, because each animal box is fingerprinted. CCT’s labels contain errors, and the species crops came from labeled boxes rather than detector boxes. SpeciesNet was trained partly on datasets from LILA BC, and in an independent comparison it was more accurate on LILA images than on other images for 75% of taxa (Clarfeld *et al*., 2025). The Oregon Critters results bear this out: SpeciesNet’s balanced accuracy there was 0.17 lower than on CCT (0.75 vs. 0.92), and the project model’s 0.10 to 0.19 lower, the gap narrowing as labels were added, although Oregon Critters’ 38 categories, against CCT’s 14, also make it a harder test. SpeciesNet’s other training images, from Wildlife Insights users, are not public, so over-lap with Oregon Critters, or with Nkhotakota Camera Traps, cannot be ruled out entirely. Both of the species evaluation’s datasets come from the western United States, and the mapping of SpeciesNet’s labels to each dataset’s categories, including the allowance for “Bird”, involves judgment; so does it in the occupancy evaluation, where counting only the narrowest category a label falls in cost SpeciesNet its related species. All timings come from one Mac laptop. A frozen feature extractor may level off below what fine-tuning can reach for look-alike species; fine-tuning the extractor itself on a GPU is a possible later option. The occupancy evaluation drew its partial reviews at random, by capture event, whereas reviewers work through lists such as the close calls; the full review is itself an estimate, from labels that contain errors; and two eight-week surveys cover few of the designs camera-trap studies use. Royle–Nichols abundance rests on assumptions about how animals are distributed and detected that camera data often break (Gilbert *et al*., 2021), which is why WildObserve reports it only as relative abundance. Finally, I have not yet measured how much faster reviewers work with suggestions than without them; a timed comparison with existing tools on real projects is a logical follow-up.

In summary, WildObserve shows that AI-assisted camera-trap review need not require uploading images, a GPU, or machine-learning expertise. A general species model gives useful suggestions from the first image, and a project model built on its fingerprints learns each project’s own species from just a few labels, retraining rapidly as the reviewer works. The reviewed records, and the suggestions on images not yet reviewed, then go straight into occupancy and abundance estimates.

## Software availability

### Name

WildObserve, version 0.6.4.

### Developer

Quantitative Science Consulting, LLC; https://www.qsc.earth.

### First available

2026.

### Hardware

a Mac with Apple silicon running macOS 14 or later, a PC running Windows 10 or 11, or a Linux computer; developed and tested on a MacBook Pro with an M1 Pro chip and 16 GB of memory. The macOS installer is 425 MB and the installed application, models included, about 570 MB.

### Software required

none beyond the installer, which bundles the models. R and unmarked only to run the exported occupancy script; this export is not required to run occupancy and abundance estimates in WildObserve.

### Internet

optional; used only for update checks, license activation, Google sign-in, and alert emails, which an administrator turns on and which go through the organization’s own mail server. An offline mode turns it off.

### Availability and cost

from the developer (https://www.qsc.earth/WildObserve). The Community edition is free and needs no license. Reviewers of this manuscript can request a free advanced license from the author.

WildObserve™ is a trademark of Quantitative Science Consulting, LLC.

## Acknowledgments

I thank the Microsoft AI for Good Lab and the MegaDetector team, and Google and the SpeciesNet team, for releasing their models under permissive licenses, and the California Institute of Technology, the Oregon Critters and Nkhotakota Camera Traps teams, African Parks, Malawi’s Department of National Parks and Wildlife, and LILA BC for making Caltech Camera Traps, Oregon Critters, and Nkhotakota Camera Traps available. I also thank the thousands of scientists and technicians who have annotated camera-trap images.

## Supporting Information

### Appendix S1: Software description

This appendix describes what WildObserve does, in the order in which a project uses it. Table S1 lists the same features for quick reference.

#### Projects and import

A project references one or more image folders, which may sit on an external drive; the images stay where they are and are never modified. WildObserve reads each image’s capture time, GPS position, and camera make, model, and serial number from its EXIF metadata. The user chooses which folder level is a deployment and which levels name places and cameras (for example, `Site/Camera in Site/Camera/Date/IMG_0001.JPG`, so that all of one camera’s memory cards form one deployment), and each deployment can be given a location for exports that need one. Because camera clocks drift or are incorrect, each deployment has a clock correction option, set from the true time of its first image, that applies wherever times are shown or exported. Each deployment can also record when its camera was set up and taken down, for the effort in occupancy models; left blank, its first and last images stand in.

Thumbnails are cached in the project, so images can be browsed while their drive is unplugged, and a moved or renamed folder can be re-linked without losing any work. Copies of an image, found by their SHA-256 checksum, are counted once, and the exports list only the original, so an archive imported twice does not double its records.

Consecutive images from one deployment are grouped into capture events, by default images no more than 60 s apart, which matches how camera traps fire in bursts. The reviewer can split an event or join it to the one before. A separate independence interval (default 30 min) groups a species’ detections at a deployment into independent records for summaries and exports (Meek *et al*., 2014).

#### Detection

WildObserve detects animals, people, and vehicles with MegaDetector v6 (Beery *et al*., 2019; Hernandez *et al*., 2024). It uses the compact variant with an Apache-licensed RT-DETRv2 architecture and a ResNet-18-vd backbone (Zhao *et al*., 2024; Lv *et al*., 2024), whose weights are published under CC-BY-4.0 (Miao *et al*., 2025); I converted it to the ONNX format. Each image is decoded at reduced resolution, which the JPEG format allows, resized to 640 *×* 640 pixels, and processed once. Every detection scoring at least 0.2 is stored. The threshold for showing a detection (range 0.2–1; default 0.5) is applied when the data are read, so users can change it at any time without reprocessing images. Each processing run records the model’s identity and SHA-256 checksum, the processor used, and the storage threshold. Processing runs in the background while the reviewer works, can be paused and resumed, and shows an estimate of the time left. An image’s detections and its “done” mark are written to the database together, so an interrupted run resumes without losing or repeating work.

Detectors often box the same stump, rock, or branch in image after image from one camera, and such boxes accounted for most of the boxes on empty images in the evaluation below. WildObserve looks for these *fixed objects*: spots in one camera’s view where boxes of nearly the same size and position (intersection-over-union of at least 0.8, each covering at most a fifth of the image) recur in at least five capture events on two or more days. It lists them for a reviewer, who confirms or dismisses each. Nothing is hidden until a person confirms it; once confirmed, boxes at that spot are left out everywhere, from the lists and counts to the exports. A reviewer can also mark a fixed object directly from a box.

#### Species suggestions

##### Fingerprints

For each of the five highest-scoring animal boxes on an image (any others when the image is opened), WildObserve crops the box, resizes it to 480 *×* 480 pixels, and runs the SpeciesNet v4.0.3a crop classifier, an EfficientNetV2-M network trained on more than 65 million camera-trap images (Tan & Le, 2021; Gadot *et al*., 2024; Google, 2026). It keeps the 1,280 numbers from the network’s final pooling layer as the box’s fingerprint and stores them once, keyed by the box’s coordinates. (EfficientNet contains many pooling layers; only the final one gives useful features, and taking an earlier one gave a balanced accuracy, defined below, of only 0.22–0.42 in development.) SpeciesNet’s final layer is a linear classifier, so WildObserve stores it separately and applies it to the stored fingerprints. This gives scores for SpeciesNet’s 2,498 labels without running the network again.

##### Project model

Once the reviewer has labeled at least three boxes each of at least two species, WildObserve trains a project model: the fingerprints are standardized and fed to a multinomial logistic regression with class-balanced weights (inverse regularization strength *C* = 0.5, fitted by L-BFGS in NumPy (Harris *et al*., 2020)). Training runs in the background: after every new label at first, then once a tenth of the labels have changed (and at least every 25 changes), straight away when a species gains enough labels to be learned, and whenever the reviewer asks; it is skipped when the labels have not changed. Boxes on images that the reviewer flagged as uncertain are left out.

WildObserve reports a cross-validated estimate of the model’s balanced accuracy (Eq. 1, in the main text). All boxes from one capture event are kept in the same cross-validation fold, so that frames of one burst never test each other. Each model version is stored as plain numeric arrays, never as executable code. Every version is kept with its balanced accuracy, so the reviewer can return to an earlier one, and any version can be exported as a model package (see Automation, below).

##### Suggestions

For each animal box, WildObserve lists the project model’s top species first, in the project’s own names, then SpeciesNet’s top labels. The project model’s top suggestion is filled in as the box’s species, so that one key press accepts it, only if the two models agree and the more confident of them scores at least 0.6. Nothing is saved until the reviewer accepts or changes it. The models agree when their names match or when SpeciesNet’s label was SpeciesNet’s top guess on at least a quarter of that species’ training boxes (for example, SpeciesNet’s “grey fox” for a project’s “gray fox”). When they disagree, nothing is filled in and the box is flagged. This guards against a weakness of any classifier trained only on the project’s labels: it can only name species it has been taught, and it may be confident even about an animal it has never seen. Before a project model exists, SpeciesNet’s top animal label is filled in when its score is at least 0.6.

#### Review

The reviewer steps through images with the keyboard (Fig. 2). Boxes can be accepted, moved, resized, deleted, or drawn, and each has a class (animal, person, or vehicle) and, for animals, a species. One species can be applied to every animal in an image, and a species list gives each of the project’s species a key, so that naming an animal takes one key press. Typed names are matched to the project’s existing spelling, and a species can be renamed or merged across the whole project. One key empties an image of boxes, to save it as empty. Images can be zoomed and panned to check small animals. An image can be flagged as uncertain, which keeps it out of the project model’s training.

Lists gather the images that need the same kind of attention, so that a reviewer can work through one kind at a time: images still to review; images with detections; likely empty images, with no box at the display threshold; images with people or vehicles; close calls, ordered by how little the top suggestion leads the runner-up, with boxes on which the two models disagree first; new species, which puts first one image of each suggested species the project has not yet labeled; images the rules flagged; reviewed images; images saved by the rules; images marked uncertain; and copies of images added earlier. Any list can be narrowed by place, camera, deployment, dates, or species, and lists stay quick to scroll at a million images because they are paged and drawn only where they are in view. From the likely-empty list, the reviewer can mark every image empty at once. By default, this skips images whose capture event has a box in another frame, because the animal may show only in that frame, and one undo restores them all.

The review screen shows the open image’s capture event as a filmstrip, frame by frame with its boxes; frames outside the current list can be opened from it, and the reviewer can split or join the event there. The animals in chosen capture events can be named at once, which suits bursts of one animal, with a warning when an event is far longer than a burst and an undo. Each box can record a count, sex, life stage, and behavior, and each image a note. For each event, WildObserve shows a tally: the largest number of each species in any one of its images, a common way to count individuals in a burst without counting the same animal twice. A dashboard summarizes review progress, what the images hold, and the species found, and timed review sessions measure how many capture events a reviewer reviews per hour, with pauses left out.

#### Automation

##### Production models

A project model can be exported as a model package: a file whose manifest lists every file’s SHA-256 checksum, the model’s license, and its training-data provenance. An administrator imports packages into a model store shared by the computer’s projects. On import, WildObserve checks the package’s layout, manifest, checksums, and license metadata, loads the model, and checks that it gives the outputs its manifest describes; a package that fails any check is refused with the reason. Each project can then choose one model as its *production model*, which scores images as they are processed, and every choice is logged. A project can thus apply a model trained on one archive to new images from the same study.

##### Watched folders

A project’s image folders can be watched, so that new images, for example from a folder synced with field computers, are added and processed as they arrive. Folders are checked every 30 s, and a file is added only once it has stopped changing for at least 10 s, so an image still being copied is not read half-written. Watching continues for projects that are not open, and after a restart. On a Mac, closing the window leaves WildObserve working from the menu bar while there is work, keeping the computer awake, and it can start at login.

##### Rules

Per-project rules act on each newly processed image. They can save confident empty images (no box scoring 0.2 or more) and confident species by themselves. Both are off by default, and records saved this way are credited to the rules, listed apart from people’s reviews, and kept out of the project model’s training, so a reviewer can check and correct them. The rules also flag images with people, species not on the project’s list, animals for which no species stands out, and species of concern; species of concern also raise a notification on the computer.

##### Alerts

An administrator can turn on alerts by email for species of concern. When a watched folder brings in an image of one, WildObserve optionally emails the species, score, place, and time, and the photo if chosen, through the organization’s own mail server, once per capture event and, by default, at most once an hour for a species at one deployment. The mail server’s password is kept in the operating system’s keychain.

#### Records and export

Detections are never edited. The reviewer’s boxes are saved as separate annotations, each linked to the detection it came from; an accepted suggestion records which model made it and its score; and each save appends the full before-and-after state of the image to an append-only review log, credited to the person who saved it.

Reviewed images, including those confirmed as empty, can be exported in eight formats: Camtrap DP 1.0.2 (Bubnicki *et al*., 2024); a flat CSV table; COCO Camera Traps and plain COCO; MegaDetector’s batch-output format, which Timelapse and AddaxAI open; YOLO labels, for training detectors; a bulk-upload package for Wildlife Insights; and JSON with full provenance. The Camtrap DP, Wildlife Insights, MegaDetector, and COCO exports pass their formats’ own validators. The Camtrap DP export records each observation as classified by a person, with the reviewer, the time of review, and the capture event. The CSV and COCO exports also record, for each box, whether the reviewer kept, adjusted, or drew it, the detector’s confidence, and whether a model suggested its species name (which model, with what score) or the reviewer typed it. Scientific names come from SpeciesNet’s taxonomy, directly or through the names the project model learned. The provenance export keeps apart, for each image, what the models said and what reviewers saved, with every save’s before-and-after history, and lists every model that touched the data by checksum: each detector run with its storage threshold, every project model version with its training labels’ checksum and cross-validated balanced accuracy, and the production models used. Detection histories for occupancy and abundance models are exported separately, as described next.

#### Occupancy and abundance

##### Detection histories

From a project’s records, WildObserve builds the detection histories that occupancy and abundance models take (Fig. 3). A site is a place, so that a camera’s successive memory cards pool together, or a single deployment. Each site’s survey is divided into occasions of a fixed number of days, seven by default, because camera-trap detections cluster in time and shorter occasions bias occupancy low (Goldstein *et al*., 2024); occasions are counted from each site’s first day or from the survey’s. A site’s effort in an occasion is its camera-days: the days on which one of its cameras ran for any part of the day, from when the camera was set up to when it was taken down, or from its first image to its last. An occasion without camera-days is missing, not a non-detection. A species is detected in an occasion when a reviewed image shows it. Optionally, an image not yet reviewed counts too, when the top suggestion for one of its animal boxes (the one the review would offer first, from the project model if there is one) names the species with a score of at least 0.9 by default. Only boxes at the display threshold count, never boxes at a confirmed fixed object, and suggestions can be limited to the species on the project’s list. Each occasion also records the most individuals of the species in any one image.

##### Models

WildObserve fits three models to each species by maximum likelihood. In single-season occupancy (MacKenzie *et al*., 2002), site *i* is occupied (*z*_*i*_ = 1) with probability *ψ*_*i*_, and if it is, the species is detected there in occasion *j* with probability *p*_*ij*_:

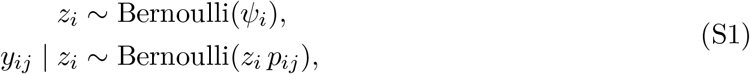

where logit *p*_*ij*_ is linear in the occasion’s camera-days when they vary. In the Royle–Nichols model (Royle & Nichols, 2003), site *i* holds *N*_*i*_ *∼* Poisson(*λ*_*i*_) animals, each detected in an occasion with probability *r*_*ij*_, so that the species is detected with probability *p*_*ij*_ = 1 − (1 − 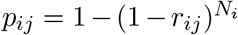 ; the likelihood sums over *N*_*i*_ up to 50. Because *λ*_*i*_ rests on the model’s assumptions about how animals are distributed and detected, WildObserve reports it as relative abundance. N-mixture models are left out, because camera-trap counts seldom meet their assumptions. When suggestions count, a third model allows for their false positives (Miller *et al*., 2011). Each occasion’s outcome is then 0 (not detected), 1 (detected by suggestions alone), or 2 (detected in a reviewed image). At an occupied site, the species is detected with probability *p*_*ij*_, and a detection is confirmed by a reviewed image with probability *b*_*ij*_; at an unoccupied site, nothing is confirmed, and suggestions alone show the species with probability *f*. The probabilities of outcomes 0, 1, and 2 are

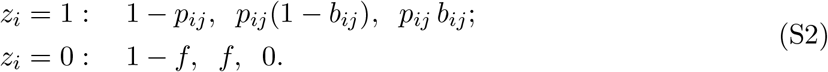

Logit *b*_*ij*_ is linear in the share of the occasion’s animal images that were reviewed: the more of them a person has checked, the more likely a true detection is to be confirmed. This is the design that unmarked’s `occuFP` fits as type 3 data (Miller *et al*., 2011; Chambert *et al*., 2015). WildObserve shows its estimate beside occupancy from reviewed images alone.

##### Covariates and fitting

Site covariates, such as habitat, elevation, or whether a camera faces a trail, are imported from a CSV file with a row per place. Numeric covariates are standardized, so that their effects are per standard deviation, and each level of a categorical covariate is compared with its first. The user chooses which covariates occupancy (or abundance) and detection depend on, each linearly on its link scale; occupancy, abundance, and detection are then reported as averages over the sites. The likelihoods are maximized in NumPy (Harris *et al*., 2020) by L-BFGS, with their analytic gradients. Standard errors come from the inverse of the Hessian, found by finite differences of the gradient, and 95% intervals are made on the link scale and transformed back, as unmarked makes them (Fiske & Chandler, 2011). No R installation is needed.

##### Estimates, plots, and export

For each species, a table gives the share of sites where it was detected and the estimates with 95% intervals: occupancy; the chance of detecting the species at an occupied site in a full occasion; relative abundance; the covariates’ effects; and, when suggestions count, occupancy from reviewed images alone, occupancy allowing for false positives, and *f*. Warnings accompany the estimates when there are fewer than 20 sites; when fewer than half of the sites had a camera running in more than one occasion; when images not yet reviewed count as non-detections, or suggestions may add false detections; when a species was detected at fewer than three sites; when a fit did not converge or an estimate reached its limit; and when sites without a value for a chosen covariate are left out. Six plots show each species’ occupancy beside the share of sites where it was detected; a species’ predicted occupancy or detection across one covariate; the covariates’ effects with their intervals; the camera-days needed for a 95% chance of detecting a species at a site it occupies; when each site’s camera ran and on which days a species was detected; and each site’s probability of being occupied, given what was seen there. Each plot can be saved as SVG or PNG. The estimates can be exported as a CSV table, and the detection histories, counts, effort, site covariates, and reviewed shares as CSV tables with an R script that fits the same models with unmarked, so that an analysis can be continued there.

#### Accounts and administration

An administrator adds reviewers and other administrators, who sign in to local accounts with a password or a Google account (Architecture, in the main text). A project can be shared across an organization’s computers. An administrator can switch WildObserve to offline mode, which turns off its four online tasks: the update check, license activation, Google sign-in, and alerts. When something goes wrong, the user can save a diagnostic report and send it to the developer; it may hold file and folder paths, but never images, species names, reviews, or account details, and WildObserve never sends it by itself.

**Table S1:** What WildObserve does, by stage of the workflow, summarizing Appendix S1.

| Feature | What it does |
| --- | --- |
| *Projects and import* |  |
| Projects | A folder holding the project’s databases; image folders stay in place, on any drive, and are re-linked if moved; copying the folder moves the project, reviews and models included. |
| Image folders | Cached thumbnails let images be browsed while their drive is unplugged; copies count once. |
| Deployments | Folder levels name deployments, places, and cameras; capture time, GPS position, and camera serial number come from each image; clock corrections; set-up and take-down times. |
| Capture events | Images no more than 60s apart at a deployment form an event, which can be split or joined; an independence interval (default 30 min) groups records for summaries and exports. |
| *Detection and species suggestions* |  |
| Detection | MegaDetector v6 finds animals, people, and vehicles in the background, with pause, time left, and resumption; boxes from 0.2 are kept, so the display threshold can change later. |
| Fixed objects | Spots that the detector boxes again and again at one camera (at least five capture events on two or more days) are found, or marked by hand, and hidden everywhere once confirmed. |
| SpeciesNet | Suggests a species for every animal from its 2,498 labels, with no project labels; until a project model exists, its top label is filled in when it scores at least 0.6. |
| *Review* |  |
| Review screen | By keyboard: boxes are accepted, moved, resized, deleted, or drawn, each with a class and species; one species for all animals; zoom and pan; uncertain images kept out of training. |
| Image lists | To review, with detections, likely empty, people and vehicles, close calls, new species, flagged, reviewed, saved by rules, uncertain, duplicates; narrowed by place, camera, deployment, dates, or species. |
| Marking empty | Likely-empty images marked empty together, skipping bursts with boxes, with undo; one key empties an image. |
| Event filmstrip | The open image’s capture event, frame by frame with boxes; animals in chosen events named at once. |
| Observations | A count, sex, life stage, and behavior per box, a note per image, and each species’ tally per event. |
| Species names | A key per species; typed names matched to the project’s spelling; renaming and merging across the project; scientific names from SpeciesNet’s taxonomy. |
| Dashboard | Review progress, what the images hold, and the species found; timed sessions measure events reviewed per hour. |
| *Species models* |  |
| Project model | Learns the project’s species from reviews, retraining by itself; balanced accuracy cross-validated by capture event; versions kept, restorable, and exportable; suggestions filled in only when SpeciesNet agrees. |
| Production models | Model packages, whose manifests list their checksums, license, and training-data provenance, are imported into a shared store; each project chooses a production model, which scores images as they are processed; every choice is logged. |
| *Automation* |  |
| Watched folders | New images are added and processed on arrival, even in closed projects; on a Mac, work continues from the menu bar, keeps the computer awake, and can start at login. |
| Production rules | Save confident empty images and species (off by default; credited to the rules, listed apart, kept out of training); flag people, unlisted species, unclear animals, and species of concern. |
| Alerts | Emails about a species of concern, with the species, score, place, and time, and the photo if chosen, through the organization’s own mail server, once per capture event. |
| *Records and export* |  |
| Review log | Model output is never edited; each save appends the image’s full before-and-after state to an append-only log, credited to the reviewer. |
| Exports | Camtrap DP 1.0.2, CSV, COCO Camera Traps, COCO, MegaDetector batch output (which Timelapse and AddaxAI open), YOLO labels, and a Wildlife Insights bulk upload. |
| Provenance | JSON with what the models said and what reviewers saved, kept apart, every save’s history, and every model that touched the data, by checksum. |
| *Occupancy and abundance* |  |
| Models | Detection histories (camera-days as effort) from reviews and optionally suggestions; occupancy, Royle–Nichols abundance, and false-positive occupancy, with site covariates. |
| Plots and export | Six plots (SVG or PNG); estimates and detection histories as CSV, with an R script for unmarked. |
| *Accounts and administration* |  |
| Accounts | Local accounts for administrators and reviewers, who sign in with a password or a Google account; every review is credited to its reviewer. |
| Shared projects | A project shared across an organization’s computers. |
| Administration | Optional internet (update checks, license activation, Google sign-in, alerts), off in offline mode; a diagnostic report the user saves and sends, without images or reviews. |

